# Molecular phylogeny and historical biogeography of andrenine bees (Hymenoptera: Andrenidae)

**DOI:** 10.1101/2020.06.09.103556

**Authors:** Gideon Pisanty, Robin Richter, Teresa Martin, Jeremy Dettman, Sophie Cardinal

**Affiliations:** Canadian National Collection of Insects, Arachnids and Nematodes, Agriculture and Agri-Food Canada, 960 Carling Ave., Ottawa, Ontario, K1A 0C6, Canada; Agriculture and Agri-Food Canada, 960 Carling Ave., Ottawa, Ontario, K1A 0C6, Canada; Steinhardt Museum of Natural History, Tel Aviv University, Tel Aviv 69978, Israel

**Keywords:** *Andrena*, Andreninae, Beringia, phylogenomics, subgeneric classification, ultraconserved element

## Abstract

The mining bee subfamily Andreninae (Hymenoptera: Andrenidae) is a widely distributed and diverse group of ground-nesting solitary bees, including numerous species known as important pollinators. Most of the species diversity of Andreninae is concentrated in the mainly Holarctic genus *Andrena*, comprising ca. 1500 described species. The subfamily and especially the genus have remained relatively neglected by recent molecular phylogenetic studies, with current classifications relying largely on morphological characters. We sampled ultraconserved element (UCE) sequences from 235 taxa, including all andrenine genera and 98 out of 104 currently recognized *Andrena* subgenera. Using 419,858 aligned nucleotide sites from 1009 UCE loci, we present a comprehensive molecular phylogenetic analysis of the subfamily. Our analysis supports the recognition of seven distinct genera in the Andreninae: *Alocandrena, Ancylandrena, Andrena, Cubiandrena, Euherbstia, Megandrena*, and *Orphana*. Within the genus *Andrena*, present-day subgeneric concepts revealed high degrees of paraphyly and polyphyly, due to heavy morphological character homoplasy, necessitating a thorough, extensive revision of the higher classification of the genus. Our results also show that the MRCA of *Andrena*+*Cubiandrena* dispersed from the New World to the Palaearctic probably during the Eocene– early Oligocene, followed by 10–14 Neogene dispersal events from the Palaearctic to the Nearctic and 1–6 Neogene dispersals back into the Palaearctic, all within the genus *Andrena*.

**Highlights:**

- A phylogeny is presented for the bee subfamily Andreninae based on UCE sequences
- The Eastern Mediterranean subgenus *Cubiandrena* is sister to all remaining *Andrena*
- Many *Andrena* subgenera exhibit paraphyly or polyphyly, requiring recircumscription
- At least 11 Old World–New World exchange events have occurred within *Andrena*

## 1. Introduction

Andreninae is a subfamily of ca. 1567 described species of ground-nesting, solitary to communal bees which is distributed throughout all continents except Australia and Antarctica (Michener 2007; Ascher & Pickering 2020), including many important pollinators of crops and wildflowers (e.g. Javorek et al. 2002; Schiestl et al. 2000). Of the six^1^ extant Andrenine genera (sensu Ascher 2004), five consist of 1–5 species each, all solitary pollen specialists inhabiting warm xeric habitats in the New World (Ascher 2004): *Alocandrena* in Peru, *Euherbstia* and *Orphana* in Chile, and *Ancylandrena* and *Megandrena* in southwestern North America. The sixth genus, *Andrena*, is the second largest genus of bees, with ca. 1556 described species (Ascher & Pickering 2020), including solitary to communal and narrowly oligolectic to broadly polylectic taxa, occupying xeric, temperate, boreal and tropical habitats (Michener 2007). This genus is distributed mostly throughout the Holarctic, with a few species reaching Mesoamerica, sub-Saharan Africa and South Asia, and one reaching Southeast Asia; its greatest diversity occurs in semiarid Mediterranean biomes (Ascher 2004; Michener 2007). Fossil Andreninae are scarce, reaching back only to the early Oligocene (32 mya) (Michez et al. 2012), and dating estimates for the subfamily and the entire Andrenidae have varied considerably (Cardinal et al. 2013, 2018; Branstetter et al. 2017a; Cardinal 2018). Although accounting for more than 7% of the worldwide diversity of bee species, the subfamily (and the Andrenidae in general) remains relatively neglected by recent molecular phylogenetic studies, especially with regard to the intrageneric classification of the large genus *Andrena*, which currently relies mostly on morphological characters.

The classification of the subfamily Andreninae has been modified several times in recent decades. Michener (1986) revised the subfamily and recognized six genera: *Alocandrena, Ancylandrena, Andrena, Euherbstia, Megandrena* and *Orphana*. However, in his subsequent treatise of bee systematics, he placed *Alocandrena* in its own subfamily, Alocandreninae, and elevated *Andrena* (*Melittoides*) to genus level (Michener 2000). In 2007, he relegated *Melittoides* back to its previous subgeneric level (Michener 2007). Ascher (2004; reviewed in Danforth et al. 2013) conducted a combined morphological and molecular phylogenetic study of the Andrenidae family. He retained *Alocandrena* within the Andreninae and *Melittoides* within *Andrena*, and divided the subfamily into two distinct tribes – Euherbstiini, comprising *Euherbstia* and *Orphana*, and Andrenini, comprising *Alocandrena, Ancylandrena, Megandrena* and *Andrena*. Dubitzky et al. (2010) conducted a morphological phylogenetic study of the genus *Andrena*, and raised the subgenus *Andrena (Cubiandrena)* to genus level. Cardinal et al. (2018) conducted an Anthophila-wide molecular phylogenetic study of the evolution of floral sonication; their findings supported Ascher’s two-tribe classification and inclusion of *Alocandrena* in the Andrenini.

Numerous morphological studies of *Andrena* have attempted to divide this immense genus into subgenera or species groups. In part due to the enormous diversity of the genus, most systematists have focused their work on limited ranges of the genus’ wide distribution; hence, a world-wide integrated view of the systematic relationships is generally lacking. Recent worldwide taxonomic treatments have recognized as many as 96–104 subgenera of *Andrena*, each comprising between 1– 100 species (Gusenleitner & Schwarz 2002; Michener 2007; Dubitzky et al. 2010; Ascher and Pickering 2020). Existing *Andrena* subgeneric keys are notoriously complicated, because to accommodate all species many subgenera are reached from two and even three different end couplets (Warncke 1968; LaBerge 1985). The North American *Andrena* fauna has been thoroughly studied and classified by LaBerge and coworkers, which have published detailed revisions of all New World subgenera and proposed phylogenetic hypotheses for the majority of them (see Dubitzky et al. 2010 for comprehensive reference list). Similarly, the Eastern Palaearctic subgenera were thoroughly revised by Tadauchi and coworkers (see Dubitzky et al. 2010). One of the most comprehensive studies on the classification of Western Palaearctic *Andrena* was conducted by Warncke (1968), who redescribed the known subgenera, and added 21 new ones. Dylewska (1987) proposed an alternative classification of Central European *Andrena*, abandoning subgeneric concepts and using species groups instead, but her classification has been largely ignored in the literature in favour of Warncke’s approach. However, comprehensive revisions and keys are lacking for nearly all subgenera in the Western Palaearctic region, amounting to an enormous diversity of bee taxa which are poorly studied.

Several subgenera of *Andrena* have been postulated as being the earliest diverging lineages within the genus, and sometimes placed as distinct genera. Lanham (1949) suggested the Nearctic subgenus *Callandrena* as one of the most “primitive” [*sic*] *Andrena* lineages. Warncke (1968) also suggested this subgenus, as well as the Mediterranean *Cubiandrena*, as closest to the genus *Megandrena*. LaBerge (1986b) suggested the Holarctic subgenera *Andrena s*.*s*. and *Notandrena* and the Nearctic *Gonandrena* as most closely resembling the most recent common ancestor of the genus. Michener (2000) considered the Mediterranean subgenus *Melittoides* as an independent lineage and raised it to generic rank. Nevertheless, rigorous studies testing the different hypotheses of relationships among *Andrena* subgenera have remained sparse. Tadauchi (1982, 1985) analysed 130 female morphological characters from 76 Japanese *Andrena* species using phenetic numerical taxonomy; his findings largely supported the existing subgeneric classification, and he did not propose any evolutionary scenarios. Dubitzky et al. (2010) analyzed 162 adult morphological characters from 102 species of 84 subgenera distributed worldwide. They identified *A*. (*Cubiandrena*) or *Ancylandrena*+*A*. (*Cubiandrena*) as sister to all remaining *Andrena*, and named three new subgenera. A few molecular phylogenetic analyses have been conducted at the genus level, each sampling only 1–2 genes from a limited number of subgenera (Ascher 2004; Dubitzky 2006; Larkin et al. 2006; Gerth et al. 2013; He et al. 2018, 2019). Ascher (2004) analysed sequences of a single nuclear gene (in addition to 89 morphological characters) from 25 mostly Nearctic *Andrena* species representing 20 subgenera. He found *A. (Callandrena) brooksi* to be sister to all remaining *Andrena* sampled, which included the subgenus *Melittoides* but not *Cubiandrena*. Larkin et al. (2006) analysed sequences of one nuclear and one mitochondrial gene from 54 Nearctic *Andrena* species representing 25 subgenera, with a focus on the subgenus *Callandrena*. They found two clades of *Callandrena* to be sister to all remaining *Andrena* sampled, and identified several other subgenera which did not form natural groups.

The current study uses nucleotide sequences to unravel the relationships within Andreninae and especially within the diverse genus *Andrena*, in order to provide the phylogenetic framework needed for a stable classification of the subfamily. Our robust reconstruction of the evolutionary history of lineages within Andreninae is based on 1009 ultraconserved element loci from 235 species, representing all andrenine genera and nearly all subgenera of *Andrena*. To provide characters that can be used to identify clades and guide future decision making on classification within the genus *Andrena*, we also mapped 162 morphological characters from Dubitzky et al. (2010) onto our proposed molecular phylogeny. Lastly, we used our phylogenetic framework to estimate divergence times and obtain insights into the biogeographic history and the timing of Old World–New World faunal exchanges within Andreninae.

## 2. Materials and Methods

### 2.1 Taxon Sampling

Our dataset included 235 species of andrenid bees (Table S1). We chose five outgroup taxa from the two other subfamilies of Andrenidae, the Oxaeinae and Panurginae: *Calliopsis andreniformis, Clavipanurgus orientalicus, Perdita bishoppi, Pseudopanurgus andrenoides*, and *Protoxaea gloriosa*. Within Andreninae, we sampled one to two species from each genus (*Alocandrena, Ancylandrena, Euherbstia, Megandrena* and *Orphana*) except *Andrena*. Within the latter genus, we sampled 1–11 species from each of 98 currently recognized subgenera, depending on subgenus size and morphological diversity. From each subgenus, we tried to sample the type species or the closest available relative, and additional species that deviate markedly from the type species or belong to different species groups. We additionally sampled a few poorly classified or undescribed species, that appeared to represent independent intrageneric clades in preliminary analyses. Sequences from six monotypic *Andrena* subgenera could not be obtained for the current study: *Aporandrena, Carinandrena, Celetandrena, Chaulandrena, Malayapis*, and *Zophandrena*. For all Holarctic subgenera except *Trachandrena*, we included members from both sides of the Atlantic. All specimens were reexamined by the first author to ensure their correct identification. For species that had COI barcode sequences publicly available on GenBank or BOLD, we also extracted the barcode sequences from the background, untargeted sequencing data and compared them with the published records. All sampled specimens along with their collected location, collector, species identifier, depository, and sex are listed in Table S1.

### 2.2 Library preparation, UCE enrichment and sequencing

DNA was extracted from whole soaked specimens and/or 1–3 pulverized legs incubated in CTAB extraction buffer, and purified using phenol-chloroform extraction followed by ethanol precipitation. DNA concentration and quality were assessed using Qubit 3.0 (Thermo Fischer Scientific, Waltham, MA) and Labchip GX Touch 24 (Perkin Elmer, Waltham, MA). Some DNA extracts that showed high levels of degradation were size-selected using Select-a-Size column or bead kits (ZYMO, Irvine, CA). High molecular weight, genomic-quality extracts were sonicated using Covaris M220 (Woburn, MA, USA) to a size of ca. 550 bp. We prepared libraries with Kapa (Kapa Biosystems Inc., Wilmington, MA) or NEBNext DNA Ultra II FS (New England BioLabs, Ipswich, MA) library preparation kits, using two unique sequence tags. We purified reactions following PCR using 1.0X AMPure substitute. For UCE enrichment we pooled 10–14 libraries together at equimolar concentrations and adjusted pool concentrations to 147 ng/µl. For each enrichment we used a total of 500 ng of DNA (3.4 µl each pool), and we performed enrichments using the V2-Hymenoptera bait set which includes 31,829 probes, targeting 2,590 UCE loci (Branstetter et al. 2017b). RNA baits were hybridized with sequencing libraries at 65°C for a period of 24 hours. Enrichment levels were assessed using qPCR by comparing amplification profiles of unenriched vs. enriched pools with PCR primers matching 5–7 different UCE loci. Successfully enriched library pools were quantified by qPCR and combined at equimolar ratios in groups of 4 for sequencing. These pools were then sequenced on a MiSeq (Illumina Inc., San Diego, CA).

### 2.3 Bioinformatics and Matrix preparation

Raw Illumina data was demultiplexed and converted from BCL to FASTQ format. We performed all initial bioinformatics steps using the PHYLUCE v. 1.4 software package (Faircloth 2015) and associated programs. We cleaned and trimmed raw reads using ILLUMIPROCESSOR (Faircloth 2013) and assembled contigs de novo using ABySS 1.3.7 (Simpson et al. 2009). We used PHYLUCE scripts to identify and extract UCE contigs, remove potential paralogs, align the UCE loci using MAFFT (Katoh & Standley 2013), and end-trim the alignments.

We created five alignment sets for phylogenetic analysis: 1) in the **232T-F75** set, we included 232 taxa, thus eliminating three taxa that received few captured UCE loci and whose inclusion destabilized preliminary trees (*Andrena erberi, A. halictoides* and *A. monacha*). We filtered the alignments for 75% taxon occupancy (percentage taxa required to be present in a given locus). This set contains 1009 loci and 419,858 nucleotide positions, of which 13% are gaps and 27% non-gap undetermined characters. This was the main set used in most of our analyses. 2) The **232T-F50** set used the same taxa as the previous one, but filtered the alignments for 50% taxon occupancy. 3) The **235T-F50** set was created to place the three previously excluded taxa in the Andreninae tree. It consists of all 235 taxa, with data from two specimens of *A. erberi* concatenated and analysed as a single unit, due to low UCE capture; the set is filtered for 50% taxon occupancy. 4) The **232T-F75-1004** set was created to control for potential base composition bias. It consists of the 232T-F75 set excluding five loci that deviate significantly from base composition heterogeneity, as assessed using BaCoCa (Kück & Struck 2014) (χ^2^ test, *p*≤0.05). 5) The **231T-F75-RY** was also created to control for potential base composition bias. It consists of the 232T-F75 set with *Andrena schlettereri* excluded (as it greatly destabilized the subsequent analysis, probably due to very low UCE capture), and converted to RY coding.

### 2.4 Phylogenetic analysis

We inferred phylogenetic trees using Maximum Likelihood (ML), Bayesian Inference (BI) and Species Tree (ST) methods. ML analysis was carried out on the concatenated 232T-F75 set using the best-tree plus rapid bootstrapping search (“-f a” option) in RAxML (Stamatakis 2014) with 100 bootstrap replicates, GTR+Γ model of sequence evolution. We compared three different partitioning schemes: 1) unpartitioned, 2) partitioned by locus, and 3) partitioned with the rcluster algorithm in PartitionFinder v2 (Lanfear et al. 2016), based on data pre-partitioned by locus. Similar analyses were carried out on the 232T-F50, 235T-F50, 232T-F75-1004 and 231T-F75-RY sets, using only the unpartitioned scheme.

BI analysis was carried out on the concatenated 232T-F75 set in ExaBayes (Aberer et al. 2014), using the rcluster partitioning scheme. We executed two independent runs, each with four chains (one cold and three heated chains). We linked branch lengths across partitions, and ran each search for one million generations. We used the program TRACER v. 1.6 (Bouckaert et al. 2014) to assess burn-in, convergence among runs, and run performance. We computed extended majority-rule consensus trees using the *consense* utility in ExaBayes.

For the ST analysis, we used RAxML to generate individual gene trees for all 1009 loci of the 232T-F75 set, employing a GTR+Γ model of sequence evolution. We summarized the trees in ASTRAL-III (Zhang et al. 2018) using the heuristic algorithm and inferred local posterior probabilities (PP; Sayyari & Mirarab, 2016) as a measure of topological support.

### 2.5 Divergence dating and biogeographic reconstruction

Divergence dating was carried out in BEAST2 v. 2.5.0. (Bouckaert et al. 2014), using an uncorrelated log-normal relaxed clock (UCLD) model (Drummond et al. 2006). Due to computational limitations of BEAST2 in analyzing large datasets, we followed Branstetter et al. 2017a and decreased computation time by inputting a fixed starting tree with all nodes constrained, turning off tree-search operators, and using only 50 UCE loci out of the 232T-F75 sequence data set, those which had the highest gene-tree bootstrap scores in RAxML (median value of all tree nodes). We concatenated the loci and analyzed the matrix without partitioning. As a fixed starting tree, we used the best tree resulting from RAxML analysis inferred from the 232T-F75 set, using the rcluster partitioning scheme. We chose a Yule tree prior, a diffuse gamma distribution on the mean branch lengths (ucld.mean; alpha = 0.001, beta = 1000), and a clock rate starting value of 1 × 10^−9^. Other priors were left at default values. We did not use specific fossils as direct calibration priors, since the exact phylogenetic placements of andrenine fossils are unclear. Instead, we defined calibration priors on four nodes in the tree based on an analysis of the entire Anthophila by Cardinal et al. (2018; see also Cardinal 2018). We calibrated the following crown groups, all with normal distributions: Andrenidae (the tree root) (mean = 69.5, stdev = 9.4, initial value = 0), Andreninae (mean = 44.0, stdev = 4.25, initial value = 0), Andrenini (mean = 36.0, stdev = 4.85, initial value = 0), and Euherbstiini (mean = 25.5, stdev = 7.6, initial value = 0). We performed a total of four independent runs, each progressing for 200 million generations. We also performed one search with the aligned data matrix removed so that the MCMC sampled from the prior distribution defined by our calibration points only, in order to check the posterior distributions of the priors. Trace files were analyzed in TRACER v. 1.6. We discarded a burnin of 20%, and summarized a maximum clade credibility (MCC) tree using LogCombiner v2.4.3 and TreeAnnotator v2.4.3. (Bouckaert et al. 2014). The resulting tree was used for subsequent biogeographical analysis, with outgroups removed.

Probabilistic inference of ancestral range reconstruction was performed using Lagrange (Ree & Smith 2008) to infer the centre of origin of the subfamily Andreninae and to reconstruct the biogeographical history of its lineages. We recognized the following three biogeographical regions: Neotropical (A), Nearctic (B), and Palaearctic (C). Maximum range size was set to two and the possible ranges were A, AB, B, BC and C. We coded all terminals unambiguously (except *Andrena barbilabris* which has a Holarctic distribution), based on the distributions of sampled species only (Fig. 1). We implemented only one time period, thus with dispersal probabilities constant throughout the tree, and scaled dispersal probabilities between adjacent regions by a value of 1.0.

**Figure 1.**
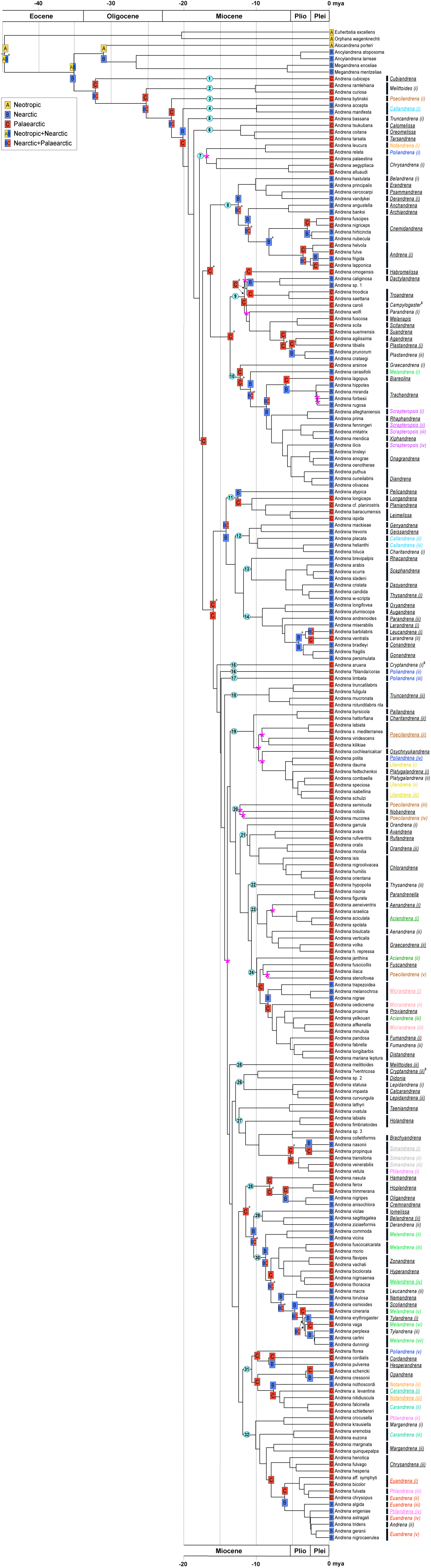
Dated Phylogeny and Biogeography of Andreninae. MCC tree inferred from a ML analysis of the 232-F75 matrix (1009 UCE loci, 419,858 bp) in RAxML, partitioned using the rcluster algorithm, followed by dating in BEAST2 using the 50 best loci. Pink asterisks indicate weakly supported nodes (<70BS), and nodes that are conflicting among ML and BI analyses performed under different partition or matrix completeness regimes or by excluding heterogeneous loci. Letters in color-coded squares at the nodes indicate ancestral ranges according to the most likely reconstruction of the Lagrange model. A black asterisk indicates that other states could not be excluded within 1.5 log-likelihood units of the maximum. Major lineages within the genus *Andrena* are numbered 1–32 in light blue circles. Black bars on the right hand side indicate current subgeneric classification of the genus *Andrena*. Each subgenus represented by three or more independent clades is notated in a different color; clades representing (or assumed to represent) the subgeneric type species are underlined. ^‡^ indicates subgenera with additional lineages, absent from the figure (see Fig. S1g).

### 2.6. Morphological character reconstruction

To find potential morphological synapomorphies that could be used to define and recognize our proposed clades, we mapped 162 characters from the external morphological dataset of Dubitzky et al. (2010) onto our inferred tree. We applied Dubitzky et al.’s character codings to the 80 species that were present also in our 232T-F75 taxon set. We additionally coded 17 species based on closely related taxa in Dubitzky et al. (2010), relying on current morphological and genetic knowledge. We further examined specimens and coded selected characters for four species representing early diverging lineages that were absent in the matrix of Dubitzky et al.: *Alocandrena porteri, Andrena bassana, A. bytinskii*, and *A. ramlehiana*. All other species were coded as missing data. Data of four characters (65, 66, 89, 94) were modified in two species already present in the original matrix (*Andrena melittoides, Euherbstia excellens*) that were examined for comparison. Our updated character matrix is given in Table S2. We mapped the characters onto our preferred phylogenetic tree (inferred using Maximum Likelihood with rcluster partitioning scheme; Fig. 1), and visualized character state changes across nodes using the program WinClada v.1.00.08 (Nixon, 2002). We optimized all unambiguous character state changes under parsimony, using either fast (ACCTRAN) or slow (DELTRAN) optimization.

## 3. Results

### 3.1 Phylogenetic analysis

Analysis of the concatenated 232-taxon matrix using the ML and BI inference methods under different gene partition and matrix completeness regimes, recovered a highly resolved and congruent phylogeny for the Andreninae with most nodes displaying high support values and only eight nodes differing among individual analyses (Figs. 1, S1a–d, h). In the ML trees (excluding the RY-encoded matrix), less than 30 nodes had BS<90, and less than 20 nodes had BS<70 (Fig. S1a–d). The BI tree was especially well resolved with all but 7 nodes with a PP value of 1.0, and 0 nodes with PP<0.6 (Fig. S1h). Only 5 loci had heterogeneous base composition, and their removal (the 232T-F75-1004 matrix) hardly affected the tree topology and support values (Fig. S1e). On the other hand, conversion of the concatenated matrix to RY coding (the 232T-F75-RY matrix) yielded a somewhat different and more weakly resolved topology, with 46 nodes having BS<90 (Fig. S1f).

Tree inference using the species tree (ST) method yielded a topology that was significantly different from the concatenated analyses, with 57 nodes that were not recovered in any of the ML and BI trees (Fig. S1i). Support values for the ST tree were relatively low, with 47 nodes having LPP<0.5. Due to their higher congruence and support values, we choose the concatenated matrix analyses (ML and BI) as our preferred analyses, but we occasionally refer also to the results of the gene tree analysis, especially when they conform better to current morphological knowledge or biogeographic patterns, and when there are incongruences among different ML and BI analyses.

### 3.2. Phylogenetic relationships within Andreninae

All our analyses recovered Andreninae, Andrenini and Euherbstiini as monophyletic groups with maximal support values, with *Alocandrena* nested in the Andrenini (Figs. 1, S1a–i). In most ML and BI analyses, the Andrenini were divided into two sister clades, *Alocandrena* + (*Ancylandrena* + *Megandrena)*, and a monophyletic *Andrena* (Figs. 1, S1a–e, h). However, in the ML analysis of the RY-encoded matrix, *Alocandrena* was nested within *Andrena* (Fig. S1f), and in the ST analysis, *Alocandrena* was sister to *Andrena* (Fig. S1i), possibly due to long-branch attraction between *Alocandrena* and *Andrena cubiceps*, whose apical branches were 2–3.5 times longer than their sister lineages (Fig. S1).

Within the genus *Andrena*, out of 51 subgenera that were sampled with at least two species, 35 were revealed as paraphyletic or polyphyletic (Fig. 1). Only three of the fourteen largest (≥30 spp.) subgenera were monophyletic: *Chlorandrena, Cnemidandrena* and *Trachandrena*. Subgenera *Poliandrena, Poecilandrena* and *Ptilandrena* exhibited the most extreme polyphyly, each represented by 4–5 independent lineages. Of the ten monotypic *Andrena* subgenera sampled, nine formed distinct lineages that were independent of other subgenera; only *Xiphandrena* was nested within another subgenus (*Scrapteropsis*) (Fig. 1). To facilitate the reporting of results and discussion, we hereby divide *Andrena* into the following 32 clades (all clades are well-supported in ML and BI analyses unless otherwise indicated):

**Clade 1**. *Cubiandrena*.

**Clade 2**. *Melittoides* (in part).

**Clade 3**. *Poecilandrena* (in part).

**Clade 4**. *Callandrena* (in part).

**Clade 5**. *Truncandrena* (in part).

**Clade 6**. *Calomelissa, Oreomelissa* and *Tarsandrena*.

**Clade 7**. *Chry*s*andrena* (in part), *Notandrena* (in part) and *Poliandrena* (in part). In the analysis of the 50% matrix, this group is split into two separate clades, branching off one after the other (Fig. S1d).

**Clade 8**. *Anchandrena, Andrena s*.*s*. (in part), *Archiandrena, Belandrena* (in part), *Cnemidandrena. Derandrena* (in part), *Erandrena* and *Psammandrena*.

**Clade 9**. *Agandrena, Campylogaster, Dactylandrena, Habromelissa, Melanapis, Parandrena* (in part), *Plastandrena, Scitandrena, Suandrena* and *Troandrena*, and an unclassified, undescribed species from Mexico (*Andrena* sp. 1). *Agandrena* is nested within *Plastandrena* (Fig. 1).

**Clade 10**. *Biareolina, Diandrena, Graecandrena* (in part), *Melandrena* (in part), *Onagrandrena, Rhaphandrena, Scrapteropsis, Trachandrena* and *Xiphandrena. Diandrena, Onagrandrena, Rhaphandrena* and *Xiphandrena* are nested within *Scrapteropsis* (Fig. 1). ML and BI analyses placed *A. ilicis* as sister to *Diandrena*+*Onagrandrena* (Figs. 1, S1a–f, h), whereas the ST analysis placed it as sister to the clade consisting of *Xiphandrena, A. fenningeri* and *A. imitatrix* (Fig. S1i).

**Clade 11**. *Leimelissa, Longandrena, Pelicandrena* and *Planiandrena*. The position of *Pelicandrena* as sister to the rest of this clade had only moderate support in some ML analyses (BS=69–97, Fig. S1a–e), and was not recovered in the ST and RY-encoded ML analyses (Fig. S1f, i).

**Clade 12**. *Geissandrena, Genyandrena, Callandrena* (in part) and *Charitandrena* (in part).

**Clade 13**. *Dasyandrena, Rhacandrena, Scaphandrena* and *Thysandrena* (in part). ST and RY-encoded ML analyses placed *Rhacandrena* in clade 14 or sister to it (Figs. 1, S1f, i).

**Clade 14**. *Augandrena, Conandrena, Gonandrena, Larandrena, Leucandrena* (in part), *Ox*y*andrena* and *Parandrena* (in part). *Conandrena, Gonandrena* and clade (i) of *Leucandrena* are all nested within *Larandrena* (Fig. 1).

**Clade 15**. *Cryptandrena* (in part).

**Clade 16**. *Poliandrena* (in part).

**Clade 17**. *Poliandrena* (in part).

**Clade 18**. *Truncandrena* (in part).

**Clade 19**. *Charitandrena* (in part), *Osychnyukandrena, Pallandrena, Platygalandrena, Poecilandrena* (in part), *Poliandrena* (in part) and *Ulandrena*. The internal topology of this clade is unstable; however, in all analyses, *Platygalandrena, Ulandrena* and clade (iv) of *Poliandrena* are nested one in the other (Figs. 1, S1). The monophyly of clade (ii) of *Poecilandrena* has weak support (BS=29–76, LPP=0.57, Fig. S1) and is not recovered in the BI analysis (Fig. S1h).

**Clade 20**. *Nobandrena* and *Poecilandrena* (in part). This clade was only weakly supported in some ML analyses (BS=28–52, Fig. S1a–e) and was not recovered in the BI, ST and RY-encoded ML analyses (Fig. S1f, h, i).

**Clade 21**. *Avandrena, Chlorandrena* and *Rufandrena*, nested within *Orandrena* (Fig. 1).

**Clade 22**. *Thysandrena* (in part).

**Clade 23**. *Aciandrena* (in part), *Aenandrena, Graecandrena* (in part), and *Parandrenella*. The position of *A. aeneiventris* is unstable; in most analyses, it is sister to clade (i) of *Aciandrena* (Figs. 1, S1a–e, h, i), whereas in the RY-encoded ML analysis, it is sister to *A. bisulcata* + clade (ii) of *Graecandrena* (Fig. S1f). In both topologies *Aenandrena* is rendered paraphyletic.

**Clade 24**. *Aciandrena* (in part), *Distandrena, Fumandrena, Fuscandrena, Micrandrena, Poecilandrena* (in part) and *Proxiandrena. Distandrena* is nested within *Fumandrena*; these as well as *Proxiandrena* and clade (iii) of *Aciandrena* are all nested within *Micrandrena* (Fig. 1).

**Clade 25**. *Melittoides* (in part).

**Clade 26**. *Calcarandrena, Cryptandrena* (in part), *Didonia*, and *Lepidandrena. Calcarandrena* is nested within *Lepidandrena* (Fig. 1). This clade was not recovered in the RY-encoded ML analysis (Fig. S1f); in the ST analysis, it includes also *A. (Melittoides) melittoides* (clade 25) (Fig. S1i).

**Clade 27**. *Brachyandrena, Holandrena, Ptilandrena* (in part), *Simandrena* and *Taeniandrena*, and an unclassified, undescribed species from Israel (*Andrena* sp. 3). Clade (i) of *Ptilandrena* is nested within *Simandrena* (Fig. 1).

**Clade 28**. *Cremnandrena, Hamandrena, Hoplandrena*, and *Oligandrena*.

**Clade 29**. *Belandrena* (in part), *Derandrena* (in part) and *Iomelissa*.

**Clade 30**. *Hyperandrena, Leucandrena* (in part), *Nemandrena, Scoliandrena, Tylandrena* and *Zonandrena*, all nested within *Melandrena* (in part) (Fig. 1). The branching order of *A. (Melandrena) cineraria* and *A. (Tylandrena) erythrogaster* is reversed in the RY-encoded ML analysis (Fig. S1f).

**Clade 31**. *Carandrena* (in part), *Cordandrena, Hesperandrena, Notandrena* (in part), *Opandrena* and *Poliandrena* (in part). Clades (i) and (ii) of *Carandrena* are interleaved with clades (ii) and (iii) of *Notandrena* (Fig. 1).

**Clade 32**. *Andrena s*.*s*. (in part), *Carandrena* (in part), *Chrysandrena* (in part), *Euandrena, Margandrena* and *Ptilandrena* (in part), all nested one in another (Fig. 1).

ML analysis of the 235T-F50 matrix, which included the three taxa that received few captured UCE loci, placed *Andrena (Cryptandrena) monacha* as sister to clades 22+23+24, *A*. (*Campylogaster*) *erberi* as sister to clade 30, and *A*. (*Stenomelissa*) *halictoides* as sister to clade 30+*A. erberi*. All these placements received low support values (BS=28–59, Fig. S1g).

### 3.3. Dating and biogeographic analyses

Results from our BEAST dating analysis and inferred ancestral ranges suggest a Neotropic or Neotropic+Nearctic origin for the Andreninae (node age 45 mya; 95%HPD 38–51), a Neotropic+Nearctic origin for the Andrenini (35 mya; 29–41), and a Holarctic origin for *Andrena* (32 mya; 27–38) (Figs. 1, S2). Different range reconstruction scenarios suggest between two to seven events of dispersal from the Nearctic to the Palaearctic, and ten to fourteen dispersal events from the Palaearctic to the Nearctic (Fig. 1; Table S3). Poorly resolved nodes leading to multiple alternative dispersal scenarios were concentrated mainly in clades 8+9+10 and 28+29+30 (Fig. 1; Table S3).

Our most likely scenario suggests with high confidence two dispersal events from the Nearctic to the Palaearctic: one late Eocene–early Miocene event at the MRCA of *Andrena* (between 32–35 Ma), and one Pliocene event at the MRCA of the node uniting clade (i) of *A*. (*Leucandrena*) and clade (ii) of *A*. (*Larandrena*) (clade 14, 3–4 Ma) (Fig. 1). Two additional late Miocene–Pliocene exchanges are suggested by this scenario, in *A*. (*Cnemidandrena*) and in clade (i) of *Andrena s*.*s*., respectively (clade 8, 3–8 and 4–8 Ma) (Fig. 1); however, these events are inferred based on very poorly reconstructed ranges in clade 8, exhibiting alternative scenarios with nearly equal likelihood (Table S3).

Our most likely scenario further suggests thirteen Middle Miocene to Pliocene (16–3 mya) dispersal events from the Palaearctic to the Nearctic, all within the genus *Andrena* (Fig. 1; Table S3). Four events have occurred at relatively deep nodes within the tree, each giving rise to diverse lineages comprising numerous subgenera: clade 8 (13–16 Ma), clade 10 (11–12 Ma), clades 11+12+13+14 (14–16 Ma), and clades 29+30 (10–11 Ma). Six later events have occurred within specific Holarctic subgenera (although these are often paraphyletic): *Euandrena*+*Ptilandrena* (clade 32, 6–8 Ma), *Micrandrena* (clade 24, 7–8 Ma), *Notandrena* (clade 31, 8–10 Ma), *Opandrena* (clade 31, 3–10 Ma), *Plastandrena* (clade 9, 3–5 Ma), and *Simandrena* (clade 27, 3–5 Ma). Three further events have given rise to the isolated Nearctic groups *Hesperandrena* (clade 31, 8–10 Ma), *Dactylandrena*+*Andrena* sp. 1 (clade 9, 11–12 Ma), and *Oligandrena*+*Cremnandrena* (clade 28, 3–6 Ma) (Fig. 1).

Our analysis also suggests two instances of long-lasting transcontinental distributions of lineages at the early-diverging nodes of the Andreninae: a North–South American distribution of the MRCAs of Andrenini and *Alocandrena*+*Ancylandrena*+*Megandrena* (45–31 mya); and a Holarctic distribution at the base of the *Andrena* backbone lineage (32–20 mya) (Fig. 1).

### 3.4 Morphological character reconstruction in *Andrena*

Most of the morphological characters mapped onto our preferred phylogenetic tree were found to be homoplasious (Figs. S3, S4). Of the few characters which exhibited non-homoplasious synapomorphies within the genus *Andrena*, three (4, 19 and 99) had a consistency index (CI) of 100. The presence of a subgenal coronet (character 4: state 1) was synapomorphic for *Andrena*. The presence of bristles on the paramandibular process (7:1,2,3) and the reduction of the mental plate (19:1) were synapomorphic for clades 2–32 under fast optimization and 4–32 under slow optimization (data on these characters is missing for clades 2 and 3). A rounded prementum with two incomplete ventrolateral ridges was synapomorphic for *Longandrena*+*Planiandrena*+*Leimelissa* (clade 11). A strongly elongate, distinctly curled anterior hair fringe of propodeal corbicula (99:1) was synapomorphic for *Simandrena*+ *Ptilandrena* (i) (clade 27).

## 4. Discussion

### 4.1 Generic classification

Our study largely supports the current tribal and generic concepts within the Andreninae, with *Alocandrena* as an additional genus in the Andrenini (Ascher 2004; Cardinal et al. 2018). The unusual morphology of *Alocandrena* within the Andreninae, which has previously led to its placement under an independent, monotypic subfamily (Michener 2007), is mirrored by the extreme branch lengths leading to this taxon in our inferred trees (Fig. S1). The monophyly of *Andrena* is confirmed, with most analyses placing it as sister to *Alocandrena*+(*Ancylandrena*+*Megandrena*). Our study also supports the generic status of *Cubiandrena*, as proposed by Dubitzky et al. (2010). *Cubiandrena* possesses 13–14 non-homoplasious autapomorphic characters that strongly distinguish it from all other members of *Andrena* (Figs. S3, S4). All of the other early-diverging clades within *Andrena* have been traditionally classified within larger paraphyletic subgenera, each containing additional members located deep within the tree; this emphasizes their great similarity to typical members of the genus. Molecular data also indicates a strong divergence of *Cubiandrena* from the rest of *Andrena*, reflected in an early split event and a disproportional branch length (Fig. S1). Recognition of *Cubiandrena* as a genus supports all the three major taxon naming criteria proposed by Vences et al. (2013): monophyly, clade stability, and phenotypic diagnosability. The removal of *Cubiandrena* from *Andrena* maintains the latter’s monophyly in all our analyses, as well as those by Dubitzky et al. The taxon is easily distinguished from *Andrena* by the unique male genitalia, with greatly reduced gonostyli; diagnostic female characters include the extremely long ocelloccipital distance, unusual facial foveae, very strongly sloping propodeum and first tergum, and unique pubescence (Dubitzky et al. 2010). Furthermore, *Cubiandrena* comprises only two species occupying a small part of the large distributional area of *Andrena*. As proposed by Dubitzky et al., we regard the subgenal coronet as the best character that defines the genus *Andrena*, one which exists only as an abnormal modification in *Cubiandrena*.

### 4.2. Phylogenetic relationships and subgeneric classification within *Andrena*

Our study corroborates many previous studies (Lanham 1949; Warncke 1968; Michener 2000; Gusenleitner & Schwarz 2002; Ascher 2004; Larkin et al. 2006; Dubitzky et al. 2010), which postulated that *Callandrena, Cubiandrena* and/or *Melittoides* constitute early-diverging lineages within *Andrena* or distinct genera. However, our results do not support the early diversion of *Andrena s*.*s*., *Gonandrena, Notandrena* or related groups, suggested by LaBerge (1986b). Furthermore, *Andrena melittoides*, the type species of subgenus *Melittoides*, was found to be unrelated to other subgeneric members and to be nested deep within the tree, contrary to Michener’s (2000) hypothesis which regarded it as a separate genus. Our analysis further revealed two Eastern Mediterranean, monospecific early-diverging *Andrena* lineages (clades 2 and 4) that have been overlooked in previous studies of the genus. The two species forming these lineages have traditionally been classified under large, unrelated subgenera, whose typical members are nested deep within the phylogeny (*A. bassana* in *Truncandrena*, Warncke 1969; *A. bytinskii* in *Poecilandrena*, Gusenleitner & Schwarz 2002). Each of these species bears some peculiar morphologies present nowhere else within the genus.

The polyphyly of subgenera such as *Callandrena* and *Melittoides*, which are represented in both early- and late-diverging *Andrena* lineages, exemplifies the problem of extreme morphological homoplasy and the difficulty in distinguishing among ancestral vs. derived character states in *Andrena*. As shown in our analysis (Figs. S3, S4), the genus is characterized by numerous reversals and homoplasies in nearly all essential morphological characters, rendering it impossible to divide into a handful of large, easily diagnosable groups. Even when there appears to be an easy morphological character that unequivocally defines a monophyletic group, this is usually limited only to one sex (e.g. female hind femoral spines in clade 21, complete propodeal corbicula in *Simandrena*+*Ptilandrena* (i)), and there is no simple way to define the group for both sexes alike. The absence of clearly defined limits between different groups of *Andrena* has led some *Andrena* systematists to erect numerous very small subgenera (especially in the New World, LaBerge 1985) for taxa that did not clearly match any previously known subgeneric concept, resulting in a highly inflated number of subgenera for the genus, criticized by Michener (2007). Another strategy, used most often in Old World taxa, was to lump poorly-fitted taxa into otherwise well-defined groups, resulting in several very large, heterogeneous and poorly-defined subgenera. Two of the most polyphyletic subgenera found in this study, *Poecilandrena* and *Poliandrena*, are clearly the result of such a lumping approach (the “wastebasket taxon” concept, see also Pisanty et al. 2018). Taxa assigned to these subgenera are lumped together mostly due to their lack of clear-cut derived structures (e.g. femoral spines, complex terminalia). An opposite scenario is seen in the polyphyletic subgenus *Ptilandrena*. Here, convergent evolution of a few derived traits, most pronounced in the males (e.g. pronotal carination, enlarged mandibles, pointed genal area), is the reason for the false grouping of completely unrelated taxa. Similar cases of convergent male characters are known in other Hymenoptera, and are often related to different aspects of behaviour such as male to male aggression (Alcock 2013).

We herein describe three different reclassification schemes that can ameliorate the current chaotic situation of higher-level *Andrena* systematics, and reconcile to varying extents the phylogenetic and morphological requirements for a stable classification: **1)** Complete renouncement of the subgeneric classification system in favor of clearly diagnosable morphological species groups that are not supported by phylogeny, as in the genera *Colletes* and *Nomada*, **2)** Preservation of most currently recognized subgenera that are supported by the phylogeny, and erection of multiple new ones to accommodate all newly identified subgeneric clades. Morphological groups of several subgenera that are similar in morphology but unrelated phylogenetically, can be informally recognized to facilitate a broad knowledge of the genus, **3)** Extensive synonymization of closely related subgenera based on the phylogeny, so that the genus is divided into a handful of large stable subgenera, which presently would be difficult to distinguish morphologically. Each subgenus would then be divided into easily identifiable species groups which would be used in keys to the genus instead of the subgenera themselves. Of the abovementioned options, we believe that establishing large subgenera that are not well-defined morphologically (option **3**) is of little taxonomic use. Therefore we would only support this option if morphological characters could be found to define such large clades. Option **2** best represents both the phylogeny and morphology of the generic subgroupings, while preserving as much as possible from the traditional classification system used in the previous half century. However, based on the phylogeny presented herein, it would require the erection of around 25 new subgenera, bringing the total number of subgenera within *Andrena* to around 125 (although several other subgenera could possibly be synonymized, as discussed in detail below). Furthermore, if future molecular data from additional sampled species shows even more extreme levels of paraphyly/polyphyly of subgeneric concepts and poorer match with morphology, renouncing phylogenetic groupings altogether (option **1**) might be the only practical solution for a stable *Andrena* classification. Meanwhile, until further detailed molecular and morphological analyses are available, we suggest to maintain the prevailing subgeneric system while making small, easy-to-implement emendations to better represent the molecular phylogeny.

We hereby suggest a few subgeneric modifications within *Andrena* to better reflect phylogenetic relationships, within the framework of option 2 outlined above. For each clade, we indicate **A)** newly identified independent lineages that do not match current subgeneric concepts, and necessitate erection of new subgenera, **B)** taxa that should be reassigned to a different existing subgenus, **C)** groups of subgenera that are closely related both phylogenetically and morphologically, and could be collapsed into a single subgenus, **D)** phylogenetic relationships that poorly match current morphological knowledge and present classificatory problems, and **E)** taxa missing from our analyses whose phylogenetic position remains elusive and necessitate further sampling. We do not make any formal nomenclatural changes at this stage, as these would be precarious without analyzing molecular data from more taxa and giving detailed subgeneric redescriptions and diagnoses, which are beyond the scope of the present study. Although we sampled nearly all *Andrena* subgenera worldwide, our results leave numerous subgeneric concepts unresolved to varying extents and require additional taxon sampling, especially in Old World species currently assigned to *Avandrena, Campylogaster, Carandrena, Didonia, Hoplandrena, Larandrena, Nobandrena, Notandrena, Oreomelissa, Parandrena, Poecilandrena, Poliandrena, Stenomelissa* and *Trachandrena*, and the unsampled, monotypic *Carinandrena* and *Malayapis*. The phylogenetic scheme for New World *Andrena* seems clearer, but additional sampling is required most of all in *Derandrena* and *Scrapteropsis*, as well as the unsampled, monotypic *Aporandrena, Celetandrena, Chaulandrena* and *Zophandrena*. We now provide a detailed morphological account of each clade of *Andrena* in our phylogeny and discuss alternative options for reclassification.

**Clade 1** represents subgenus *Cubiandrena*, which we suggest to raise to generic status (section 4.1).

**Clade 2**. This well-defined clade represents all members of subgenus *Melittoides* except the type species *A. melittoides*, which occupies clade 25 and clearly differs from all other subgeneric members in genital shape, wing venation, glossal length, and type of pilosity. This group clearly merits erection of a new subgenus.

**Clade 3** is occupied by a single peculiar species, *A. bytinskii*, which merits a new subgenus. It is characterized by two submarginal cells in the forewing, a very long flocculus, and a unique genital shape known nowhere else in *Andrena*, with greatly elongated dorsal gonocoxite lobes.

**Clade 4** corresponds to clade A of subgenus *Callandrena* in the analysis of Larkin et al. (2006), which includes the *accepta-, aureocincta-, humeralis-, manifesta*-, and *vidalesi*-groups proposed by LaBerge (1967), although only two of these groups were directly represented in our study. LaBerge did not provide a key to his proposed species groups, and there appears to be no single character that distinguishes this clade from other *Callandrena* (clade 12). However, the subgenus has to be split by some scheme in order to resolve its polyphyly.

**Clade 5** is occupied by another single peculiar species, *A. bassana*, which also merits its own subgenus. This taxon has a flat clypeus, generally coarse body sculpture, and a unique propodeal corbicula, in which the dorsal half of the corbicular surface is fully covered with very dense velvety hair, unlike any other *Andrena* species.

**Clade 6**. This clade could perhaps be united under a single subgenus, *Calomelissa*, as *Oreomelissa* was erected as an offshoot from *Calomelissa* (Hirashima & Tadauchi, 1975), and *Tarsandrena* is also related to it morphologically (Dubitzky et al. 2010). Three species of *Oreomelissa*, not sampled in the present study, possess spines on the female hind femur, and possibly represent an unrelated lineage.

**Clade 7**. *Chrysandrena* (i) clearly differs from clade (ii) of the subgenus by the much smoother body sculpture and the presence of a dorsal gonocoxite lobe, and should be assigned a separate subgenus. The morphological relationship between *Chrysandrena* (i), *Notandrena* (i) and *Poliandrena* (i) is not clear; additional taxa currently assigned to *Notandrena* and *Poliandrena* could also belong to these clades. The monophyly of clade 7 should be reexamined with additional taxon sampling.

**Clade 8**. The relationship between *Andrena* s.s., *Cnemidandrena, Anchandrena* and *Archiandrena* has been suggested by LaBerge (1980, 1985, 1986b) and Ascher (2004). However, since *Andrena* s.s. and *Cnemidandrena* are both large, well-defined subgenera, it seems unjustified to unite this group under a single subgenus, although *Anchandrena* could be easily synonymized with *Archiandrena* or vice versa. The morphological relationship between *Belandrena* (i), *Derandrena* (i), *Erandrena* and *Psammandrena* is currently unclear. *Belandrena* (i) possibly includes also *A. spheralceae*, closely related to *A. hastulata*. The scope of *Derandrena* (i) is not sufficiently clear at this point, and this subgenus warrants further sampling. Interestingly, *Belandrena* and *Derandrena* each consist of two independent lineages appearing in the very same clades (8 and 29). *Andrena* (i) likely includes all species currently assigned to *Andrena s*.*s*. except *A. cornelli* and *A. tridens*, which can be easily separated morphologically, and should be moved to *Ptilandrena* (clade 32).

**Clade 9**. The subclade consisting of *Agandrena, Melanapis, Plastandrena, Scitandrena* and *Suandrena* is characterized by a coarsely rugose propodeal triangle and a basally broadened female hind tibial spur. This group could be easily united to a form single subgenus, *Melanapis. A. caroli* is clearly unrelated to *A. (Campylogaster) erberi* and should be assigned to another subgenus (Fig. S1g). Most species of *Campylogaster* are probably related to *A. caroli* and might justify erecting a new subgenus for this group, except *A. erberi, A. iranella* and *A. skorikovi*, which share a tomentous pilosity and strongly depressed tergal marginal zones. *Parandrena* (i) could represent additional Old World species currently assigned to *Larandrena* and *Parandrena*. The inclusion of *A. chalcogastra* in *Troandena* is questionable, and merits molecular investigation. *Andrena* sp. 1, discovered as part of this study, probably merits its own new subgenus (J.L. Neff, unpublished manuscript). Further taxon sampling is needed to resolve some of the relationships within clade 9.

**Clade 10**. The relationship between *Biareolina, Diandrena, Onagrandrena, Rhaphandrena, Scrapteropsis, Trachandrena* and *Xiphandrena* has been suggested by LaBerge (1986b) and Ascher (2004). The monotypic *Xiphandrena* could be easily synonymized with *Scrapteropsis*. Apart from this simple emendation, we do not foresee an easy classificatory solution to the paraphyly of *Scrapteropsis. Graecandrena* (i) could include additional species such as *A. argyreofasciata*; *Melandrena* (i) probably includes also *A. impolita*, closely related to *A. cerasifolii*. These two groups might warrant erection of new subgenera. *Trachandrena* appears as a monophyletic, well-defined, evolutionarily young subgenus; although we did not sample any Old World *Trachandrena*, COI barcodes of the Palaearctic *A. haemorrhoa* are much closer to North American *Trachandrena* than to any other taxon (GP, unpublished results), strongly suggesting a shared origin.

**Clade 11**. *Leimelissa, Longandrena, Planiandrena*, and the unsampled *Carinandrena* are small subgenera that are related morphologically (Dubitzky et al. 2010), three of them occurring only in Central Asia; they might warrant synonymization under the name *Planiandrena*. The subgeneric placement of *A. ispida* in *Leimelissa*, previously uncertain (Gusenleitner & Schwarz 2002), is hereby confirmed.

**Clade 12**. *Callandrena* (ii) and (iii) correspond to clades B and C+D of subgenus *Callandrena* in the analysis of Larkin et al. (2006), which include the *fulvipennis-, gardineri-, helianthiformis-, krigiana-, melliventris-* and *solidaginis*-species groups, and the *aliciae-, discreta-*, and *helianthi-* groups proposed by LaBerge (1967), respectively; however, only two of these groups were directly represented in our study. These taxa should be moved to a separate subgenus, although morphological delineation from other *Callandrena* (clade 4) seems problematic. The morphological characterization of clade 12 awaits further study.

**Clade 13**. This appears to be a strictly New World clade, and Old World groups assigned to *Scaphandrena* and *Thysandrena* (Warncke 1969; Ribble 1974) are unrelated (clades 18 and 22, respectively).

**Clade 14**. This clade includes several very small subgenera, but its morphological relationships are mostly unclear. The limits of *Larandrena, Leucandrena* and *Parandrena* are still unclear, and necessitate further taxon sampling, especially of Old World taxa. It seems likely that all Old World species currently assigned to *Larandrena* are closer to *A. barbilabris* and should be moved to *Leucandrena*; *Parandrena* (ii) might represent only New World taxa, with Old World species of *Parandrena* being unrelated (e.g. clade 9). The *faceta* group of *Leucandrena* consists of a separate, unrelated lineage, closer to *Melandrena* (clade 30) (see also LaBerge 1986a).

**Clade 15** represents a single species, *A. aruana*, whose morphology differs considerably from *A. ventricosa* (the type species of *Cryptandrena)*, including the absence of spines on the female hind femur, meriting a separate subgenus. The morphologically similar species *A. monacha* belongs to a third, unrelated lineage within the polyphyletic *Cryptandrena* (Fig. S1g), and may also warrant a new subgenus.

**Clade 16**. Additional species currently assigned to the polyphyletic subgenus *Poliandrena* might be placed in this clade, which would probably merit a new subgenus.

**Clade 17**. *A. limbata* and the related, unsampled *A. toelgiana* are unique among *Poliandrena* in the tomentous pilosity, and probably merit a new subgenus.

**Clade 18** represents all subgenus *Truncandrena* except the aberrant, clearly unrelated species *A. bassana* (clade 5). *Truncandrena* is hereby reestablished as a valid subgenus, as it is unrelated to the Nearcrtic *Scaphandrena* (clade 13).

**Clade 19**. Most taxa within this clade are characterized by a basally broadened hind tibial spur, but its inner organization needs further clarification. *Poliandrena* (iv) most probably represents only the *polita* group of *Poliandrena*, whereas other members of this polyphyletic subgenus are scattered throughout the *Andrena* tree (clades 7, 16, 17, 31). *Platygalandrena, Ulandrena* and the *polita* group are morphologically similar and could be united under *Poliandrena* or *Ulandrena*, thus resolving the paraphyly of this subclade. *Pallandrena* can be synonymized with the similar subgenus *Charitandrena*, while excluding the New World *A. toluca* (clade 12). Our results reveal a narrower, more coherent concept of *Poecilandrena*, which includes only the *A. labiata* group and possibly also the *A. viridescens* group (Pisanty et al. 2018). Taxa in both groups share typical elongated gonostyli, and are easily differentiated from other species previously assigned to *Poecilandrena*.

**Clade 20**. The monophyly of this clade awaits sampling of additional species from *Poecilandrena* (outside the *A. labiata* and *A. viridescens* groups) and *Nobandrena. A. seminuda* and *A. mucorea* are clearly morphologically distinct from both subgenera. The monophyly of *Nobandrena* is questionable, and unsampled species such as *A. compta, A. flavobila* and *A. probata* may represent an unrelated lineage.

**Clade 21** is well-characterized by spines on the female hind femur, and could potentially be unified under a single subgenus *Chlorandena*. However, this trait is missing in some species of *Avandrena* not sampled in this study, whose phylogenetic placement should be investigated.

**Clade 22**. This clade probably represents all four Old World species currently assigned to *Thysandrena*, and would thus merit a new subgenus. However, *A. ranunculorum* might be unrelated and should also be sampled.

**Clade 23**. The subgeneric placement of *A. israelica* in *Aciandrena* and *A. verticalis* in *Graecandrena*, previously considered uncertain (Gusenleitner & Schwarz 2002; Pisanty et al. 2016), is hereby confirmed. *A. bisulcata* would probably need to be assigned a new subgenus, in order to resolve the paraphyly of *Aenandrena*; *Aenandrena* (i) includes also *A. bonasia*, which strongly resembles *A. aeneiventris*.

**Clade 24**. *A. stenofovea* is very similar to *A. (Fuscandrena) fuscicollis*; *A. iliaca* also shares the latter’s fine sculpturing, weak metallic luster, yellow male clypeus and paraocular areas, and large, unwrinkled propodeal triangle; both species should be moved to *Fuscandrena*. Our phylogeny supports a broad concept of subgenus *Micrandrena* which includes the groups known as *Distandrena, Fumandrena* and *Proxiandrena*, as well as *A. yelkouan*. Most taxa included in *Micrandrena* under this concept share a stronger body sculpturing (most pronounced in the propodeal triangle) compared to other small-bodied *Andrena* (especially *Aciandrena, Fuscandrena* and *Graecandrena). Fumandrena* is especially difficult to distinguish from *Micrandrena* (Gusenleitner & Schwarz 2002), and the relatedness among these subgenera is hereby confirmed. The unusual, protuberant clypeus of *A. janthina* clearly differs from typical *Aciandrena* and probably warrants erection of a new subgenus.

**Clade 25**. This clade represents the unusual subgenus *Meltittoides* with its type species only. All other members of this subgenus are placed in clade 2. The morphological uniqueness of *A. melittoides* (slender stigma, long flagellomere 1, unusual first tergum and terminalia) has led to its classification as a distinct genus by Michener (2000). However, our analysis reveals that it is a highly derived late offshoot within *Andrena*.

**Clade 26**. *Cryptandrena* should include only *A. brumanensis, A. rotundata and A. ventricosa*, as other species are unrelated (see clade 15). The verified phylogenetic placement of the problematic subgenus *Didonia* awaits sampling of its type species, *A. mucida*. The recently erected *Calcarandrena* (Dubitzky et al. 2010) should be collapsed back into *Lepidandrena*, from which it was originally split.

**Clade 27**. *A. vetula* should be moved to *Simandrena*, as its female possesses a complete propodeal corbicula characteristic of the subgenus, and very closely resembles *A. venerabilis. Andrena* sp. 3, discovered as part of this study, possibly merits a new subgenus.

**Clade 28**. The monotypic subgenus *Cremnandrena* could possibly be synonymized with *Oligandrena*, which is morphologically related (Lanham 1949; Dubitzky et al. 2010). The monophyly of *Hoplandrena* is questionable, and further sampling of species such as *A. bucephala, A. mordax* and *A. najadana* might reveal a second lineage within this group (see also clade 31).

**Clade 29**. *Belandrena* (ii) probably includes all species currently assigned to the subgenus except *A. hastulata* and the related *A. spheralceae*, whereas *Derandrena* (ii) would include only the atypical *A. ziziaeformis*. The morphological characteristics of clade 29 are currently unclear to us.

**Clade 30**. A close relationship among the different subgenera found within this clade has been previously suggested by Ascher (2004). Contrary to Ascher’s view, however, only the *faceta* group of *Leucandrena* is assigned to this clade, whereas typical *Leucandrena* are in clade 14 (see also LaBerge 1986a). Given the high morphological homogeneity among members of *Melandrena*, and their great similarity to *Hyperandrena, Tylandrena* and *Zonandrena*, it seems justified to synonymize all subgenera within this clade (except *Leucandrena)* under *Melandrena*.

**Clade 31**. *Poliandrena* (v) probably merits a new subgenus; this group might include additional species currently assigned to *Poliandrena* and *Hoplandrena* (e.g. *A. mollissima, A. mordax, A. najadana*), and further taxon sampling is needed. *Cordandrena* and *Poliandrena* (v) are extremely distinct morphologically, and their relationship is currently unclear. *Opandrena* is hereby reestablished as a valid subgenus, as it is clearly unrelated to *Holandrena* (clade 27; Dubitzky et al. 2010). *Carandrena* should probably be synonymized with the similar subgenus *Notandrena*, as the phylogeny does not demarcate a clear border between the two groups; both subgenera require additional taxon sampling to better delineate subgeneric concepts.

**Clade 32**. The group consisting of *A. crocusella, A. elsei, A. grossella* and *A. krausiella* probably merits a new subgenus. Another subgenus could be erected for *A. euzona* and *A. eremobia*. Three species of *Chrysandrena* with smooth body sculpture and a dorsal gonocoxite lobe should be moved to a separate subgenus (clade 7). *Euandrena* should be synonymized with *Ptilandrena*, and *A*. (*Andrena s*.*s*.) *tridens* and the related *A. cornelli* should be moved to *Ptilandrena*.

### 4.3. Biogeography and life history

Our biogeographic reconstruction indicates a New World origin for the Andreninae with exchanges between North and South America. The MRCA of *Andrena* dispersed into the Old World, followed by numerous additional Old World–New World exchanges in diverse *Andrena* lineages.

Early-diverging Andrenine genera exhibit an amphitropical distribution in the New World, a disjunct distributional pattern documented in several bee groups from various families (Michener 1979, 2007). These genera, occupying xeric habitats in North and South America, were separated during the Eocene by the Central American Seaway and a region of unsuitable tropical climate, through which they somehow dispersed. Traversal of the Caribbean Sea could have taken place through long-distance wind dispersal, possibly aided by island hopping (Michener 1979, 2007; Fuller et al. 2005). Penetration into tropical biomes could have been possible through relatively temperate mountainous patches, similar to the typical habitats of most species of *Andrena* currently known from the tropics (Michener 2007; Dubitzky et al. 2010). We consider it unlikely that the Andrenini evolved as a single lineage distributed over both North and South America as suggested by our Lagrange analysis (Fig. 1), given the strong barriers separating xeric regions on each continent.

The early-diverging lineages of *Andrena* alternate between North American and (mostly Eastern) Mediterranean distributions. The only likely land passage between these two regions from the late Eocene onward is the Bering land bridge (Sanmartín et al. 2001). The Beringia region was relatively warm during the Eocene, allowing the passage of taxa adapted to xeric habitats as typical of the early-diverging lineages of *Andrena* (Sanmartín et al. 2001). However, no early-diverging *Andrena* lineages are known from the Eastern Palaearctic which connects these regions. An alternative possible scenario is early Eocene dispersal of the MRCA of *Andrena* directly into the Western Palaearctic through a North Atlantic land bridge, as suggested for the tribe Ancylaini (Praz and Packer 2014). This hypothesis assumes a much earlier dating of the early-diverging Andrenine lineages, as suggested by some studies (Ascher 2004; Dubitzky et al. 2010; Cardinal et al. 2013). The strict Mediterranean distribution of the three most early-diverging clades of *Andrena* as well as clade 5 (and possibly 7), could suggest a more localized, Palaearctic origin for *Andrena*, rather than the more widespread Holarctic distribution suggested by our analysis (Fig. 1). A Palaearctic origin for *Andrena* was also postulated by Dubitzky et al. (2010), and is contrary to the hypotheses of LaBerge (1986b) and Ascher (2004), who suggested a New World origin for the genus based on studies that focused mainly on Nearctic taxa.

The numerous Old World–New World exchanges within *Andrena* have most probably occurred through the Bering land bridge (Sanmartín et al. 2001). The more recent, Late Miocene–Pliocene exchanges were probably limited to cold-adapted taxa, as the climate of this region had cooled down (Sanmartín et al. 2001); this is evident by the typically northern distributions of most Holarctic subgenera, with many species reaching Canada and Central Europe (Ascher 2004). Our data show a directionally asymmetric pattern of trans-Beringian dispersals within the Andreninae, with numerous dispersals from the Palaearctic to the Nearctic, but only few occurring in the other direction, as also found in bumblebees and osmiine bees (Hines 2008; Praz et al. 2008).

Our knowledge of the life history of the five early-diverging *Andrena* lineages (clades 1–5) is severely deficient, with the exception of the well studied subgenus *Callandrena* (clade 4, LaBerge 1967; Larkin et al. 2006, 2008); almost nothing is known about *A. bytinskii* and *A. bassana* (the monotypic clades 3 and 5). Species in all five lineages occur mostly in xeric habitats, and are usually active during a short period in spring–early summer (clades 1–3, 5) or autumn (clade 4). Pollen host preferences are known only for *A. cubiceps* (clade 1) and members of *Callandrena*, both pollen specialists (Larkin et al. 2008; Roberts & Meulemeester 2015). However, the unusual propodeal corbicula of *A. bassana* could also suggest adaptation to collect pollen from a specific plant taxon, and the shortening of the glossa in the node uniting clades 5–32 (character 15 in Figs. S3 & S4, showing numerous reversals in more terminal nodes) might reflect a shift to a more generalist foraging strategy in these more recent lineages. Xeric distribution, pollen specialization and univoltine spring activity also characterize all other Andrenine genera as far as is currently known (except the long activity period in *Alocandrena*, Ascher 2004), strongly suggesting that these traits are ancestral within *Andrena* as well as Andreninae as a whole. The immense diversification and radiative expansion in younger clades of *Andrena* (clades 6–32) involved adaptation to cooler and/or wetter habitats in temperate, boreal and tropical biomes (e.g. in clades 6, 8, 10, 30, 32); numerous shifts in pollen hosts and breadths, with appearance of polylectic taxa (Larkin et al. 2008); numerous shifts in phenology and seasonality, with appearance of multivoltine taxa (Larkin et al. 2008); appearance of communally nesting taxa (e.g. in *Plastandrena* and *Hoplandrena*, clades 9 and 28; Michener 2007); and multiple dispersals between North America and Eurasia through the Bering land bridge (Fig. 1), as well as southbound dispersals into the Neotropic, Afrotropic and Oriental regions.

## 5. Conclusions

The current study is an important step towards a stable classification of the Andreninae, and a deeper understanding of the complex evolutionary history giving rise to the huge diversity within the genus *Andrena*. The high degree of morphological character ambiguity and homoplasy found among *Andrena* lineages, corroborate previous studies that showed a limited ability of traditionally used morphological characters to provide a reliable phylogenetic signal in bees (Rightmyer et al. 2013; Litman et al. 2016; Trunz et al. 2016; Dorchin et al. 2018). Our study revealed numerous *Andrena* lineages that do not fit current subgeneric concepts, and will require either the erection of new subgenera or a drastic revision of the present classification. Future studies should reexamine morphological traits within *Andrena*, to reveal characters that would define monophyletic lineages based on extensive taxon sampling and molecular and morphological data. These morphological traits will be used to diagnose the distinct lineages and formulate efficient, revised subgeneric keys. Such studies could improve the cluttered higher-level classification of the genus, potentially reducing the number of subgenera without compromising their diagnosability. If, however, further sampling reveals that morphological and phylogenetic concepts in *Andrena* cannot be reasonably reconciled, an alternative classification method, based solely on morphological species groups, would have to be developed anew.

The resolved phylogenetic relationships among Andreninae lineages are important for understanding the number and timing of adaptive changes in the diverse pollen host preferences of the genus *Andrena* (Larkin et al. 2008). The specific pattern and direction of shifts between oligolecty and polylecty can help to uncover the evolutionary mechanisms of host plant preference in bees (Sipes & Tepedino 2005; Larkin et al. 2008; Michez et al. 2008; Sedivy et al. 2008) – a key factor affecting bee habitat preference, persistence in anthropogenic habitats, susceptibility to decline and propensity to pollinate crops (Winfree et al. 2011; Bartomeus et al. 2013; Pisanty & Mandelik 2015).

Several poorly-studied, isolated taxa were found to form independent and even early-diverging lineages within *Andrena*, highlighting the need for generous taxon sampling based on extensive knowledge of the morphological diversity in such a speciose genus. Further taxon and gene sampling is still needed to resolve conflicting topologies in our existing tree, and assess the position of several subgenera and species groups of *Andrena* absent from our analysis. Although we generously sampled the two main *Andrena* diversity hotspots on each side of the Atlantic (the Mediterranean Basin and Western USA), the more poorly-studied *Andrena* faunas of the Neotropic, Afrotropic and Oriental regions were completely absent from our sampling, and those from the Eastern Palaearctic were very sparsely represented. Adequate sampling of diverse taxa from all these regions is important to obtain a fuller picture of the higher organization and biogeographical history of the genus, and to potentially identify additional early-diverging or independent lineages of *Andrena*. In particular, the pattern and timing of *Andrena* expansion into tropical habitats could shed further light on the function of tropical regions as barriers for the spread of xeric-adapted bee lineages. Detailed molecular genetic studies of *Andrena* are required also at subgeneric levels, to better understand the relationships within each of the numerous subdivisions of the genus. Lastly, much research effort is needed to solve the phylogenetic relationships and biogeographical history within the two remaining subfamilies of the Andrenidae (Panurginae and Oxaeinae), a family largely neglected by recent molecular phylogenetic studies (Danforth et al. 2013).

## Acknowledgements

We thank Julie Chapados, Kasia Dadej, Alex Hanright and Wayne McGregor for help in lab work and protocols, and Jackson Eyres, Brant Faircloth, Siavash Mirarab and Richard Ree for assistance with computer analyses and phylogenetic software. We thank all the generous collectors and curators who provided specimens for this study: Netta Dorchin (Steinhardt Museum of Natural History), Sam Droege (USGS), Fritz Gusenleitner (Biologiezentrum Linz), Ken-Lou J. Hung (University of Toronto), Thomas McElrath (Illinois Natural History Survey), Denis Michez and Thomas J. Wood (Mons University), John L. Neff (Central Texas Melittological Institute), Peter Oboyski (Essig Museum of Entomology), Laurence Packer (York University), Jerome G. Rozen and Corey Smith (American Museum of Natural History), Erwin Scheuchl (Ergolding, Germany), Stefan Schmidt (Zoologische Staatssammlung München), Jakub Straka (Charles University), Osamu Tadauchi and Toshiharu Mita (Kyushu University), Bernard Vaissière (INRA), and Doug Yanega (University of California Riverside). John L. Neff examined two specimens of *Andrena* sp. 1 and confirmed its status as an undescribed taxon. Erwin Scheuchl provided valuable comments on *Andrena* classification, and highlighted numerous instances of problematic subgeneric concepts. John S. Ascher and Christophe J. Praz provided helpful comments on an earlier version of this manuscript.

## GenBank Accession Number

PRJNA637821.

## Supplementary data

**Fig. S1**. Alternative Andreninae phylogenies inferred using **(a–g)** Maximum Likelihood, **(h)** Bayesian Inference, and **(i)** Species Tree methods. **a)** RAxML, 232T-F75, no partition; **b)** RAxML, 232T-F75, rcluster; **c)** RAxML, 232T-F75, partition by gene; **d)** RAxML, 232T-F50, no partition; **e)** RAxML, 232T-F75-1004, no partition; **f)** RAxML, 231T-F75-RY, no partition; **g)** RAxML, 235T-F50, no partition; **h)** ExaBayes, 232T-F75, rcluster; **i)** ASTRALIII, 232T-F75. Node labels indicate bootstrap support **(a–g)**, posterior probabilities **(h)**, and local posterior probabilities **(i)**, respectively. The positions of *Andrena erberi, A. halictoides* and *A. monacha* are highlighted in blue **(g)**.

**Fig. S2**. MCC tree of the Andreninae dated in BEAST2 based on the 50 best loci. A fixed tree topology was used, based on RAxML analysis of the 232-F75 matrix under the rcluster partitioning scheme. Horizontal bars indicate 95% HPD intervals for node datings, and node labels indicate node ages.

**Fig. S3**. The tree of our preferred molecular phylogenetic model with morphological characters of 97 taxa from Dubitzky et al. (2010) mapped on branches, using fast optimization (ACCTRAN). The phylogenetic position of 17 species (noted in parentheses) from the Dubitzky et al. data is assumed based on closely related taxa sampled in our dataset. The remaining 135 taxa for which all characters were left blank, are shown in grey. Black squares indicate non-homoplasious changes, white squares indicate homoplasious changes. Clade numbering (light blue circles) corresponds to Fig. 1.

**Fig. S4**. The tree of our preferred molecular phylogenetic model with morphological characters of 97 taxa from Dubitzky et al. (2010) mapped on branches, using slow optimization (DELTRAN). The phylogenetic position of 17 species (noted in parentheses) from the Dubitzky et al. data is assumed based on closely related taxa sampled in our dataset. The remaining 135 taxa for which all characters were left blank, are shown in grey. Black squares indicate non-homoplasious changes, white squares indicate homoplasious changes. Clade numbering (light blue circles) corresponds to Fig. 1.

**Table S1**. Collection data and UCE capture statistics for taxa included in the molecular phylogenetic analyses.

**Table S2**. Morphological data matrix based on Dubitzky et al. (2010). Taxa substituted or added to Dubitzky et al.’s original matrix and recoded characters are highlighted.

**Table S3**. Alternative ancestral ranges inferred by the Lagrange model for Andreninae nodes.

## Footnotes

1 For simplicity, unless otherwise indicated, *Cubiandrena* is treated throughout this paper as a subgenus within *Andrena*.

