## Supplementary figures and images for "Molecular phylogeny and historical biogeography of andrenine bees (Hymenoptera: Andrenidae)"

### Figure S1

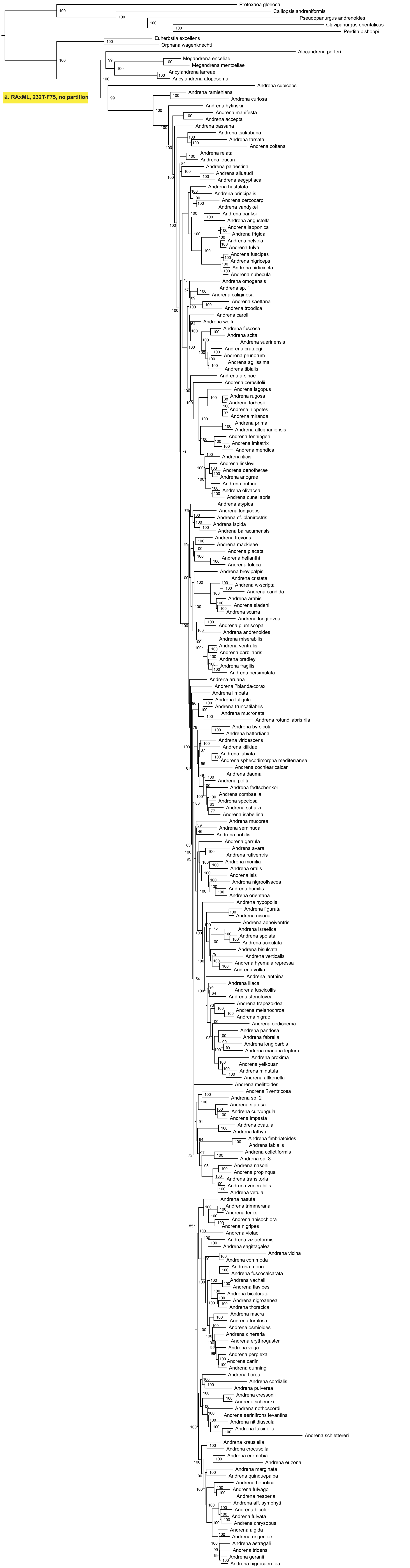

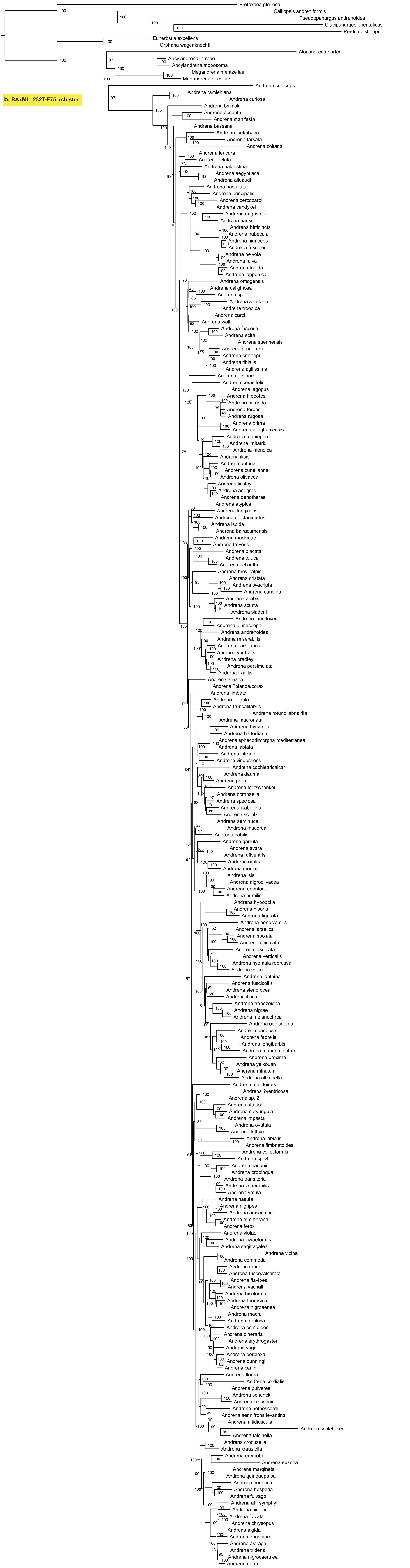

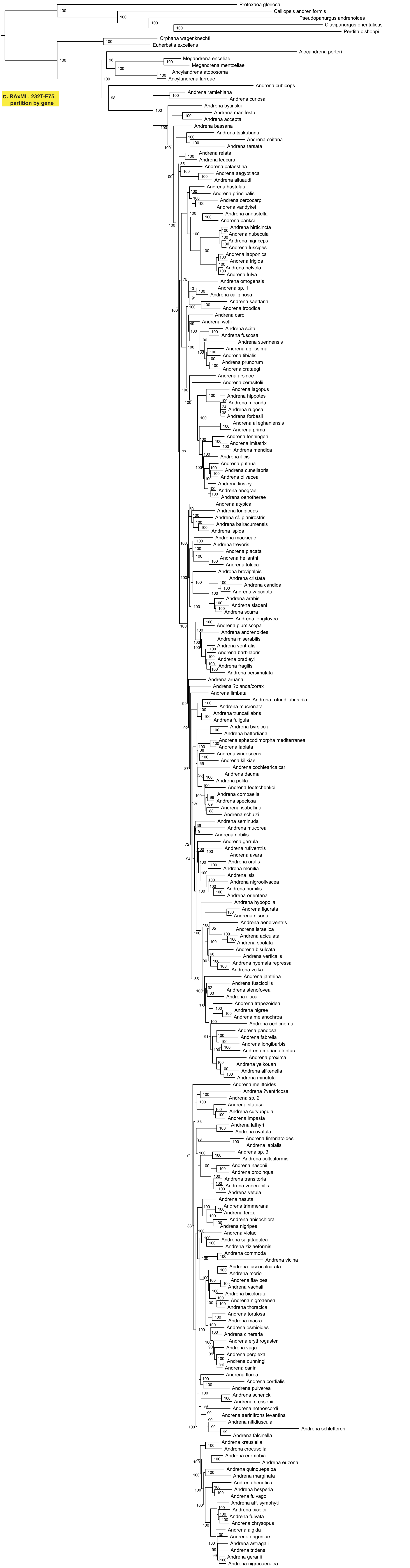

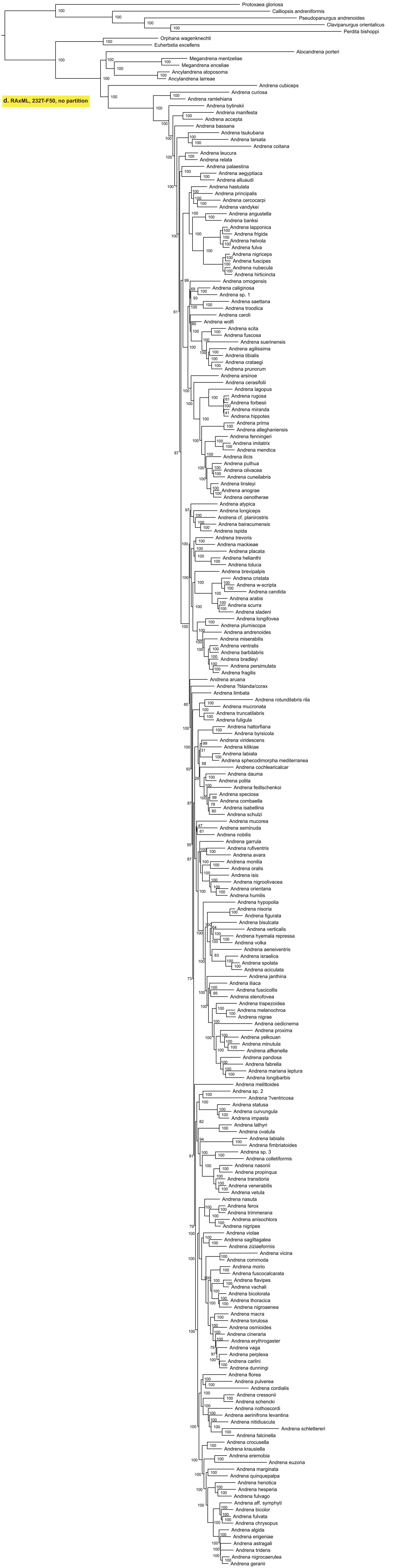

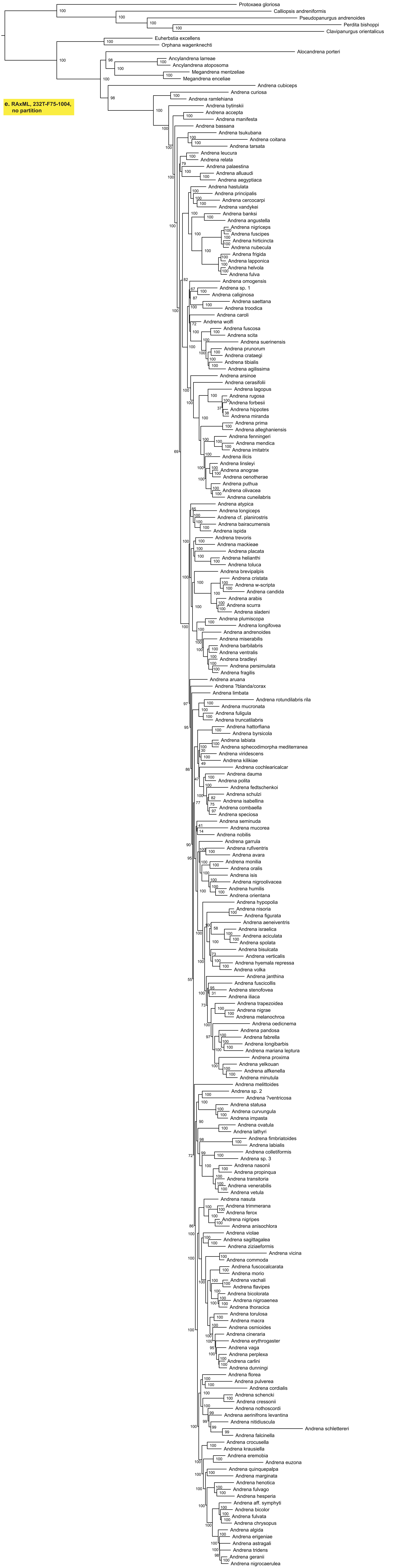

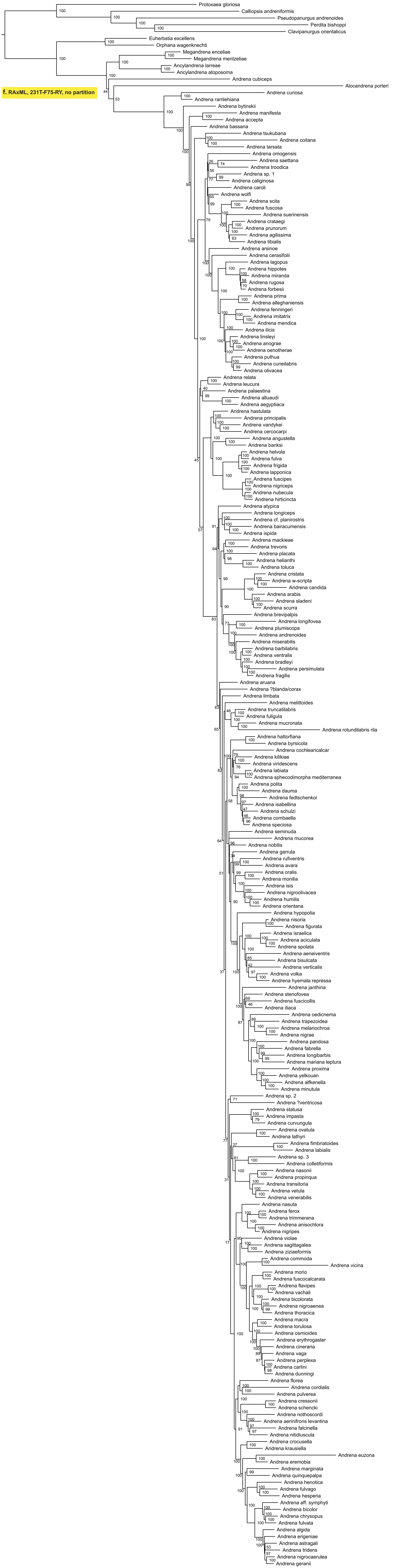


**f. RAxML, 231T-F75-RY, no partition**

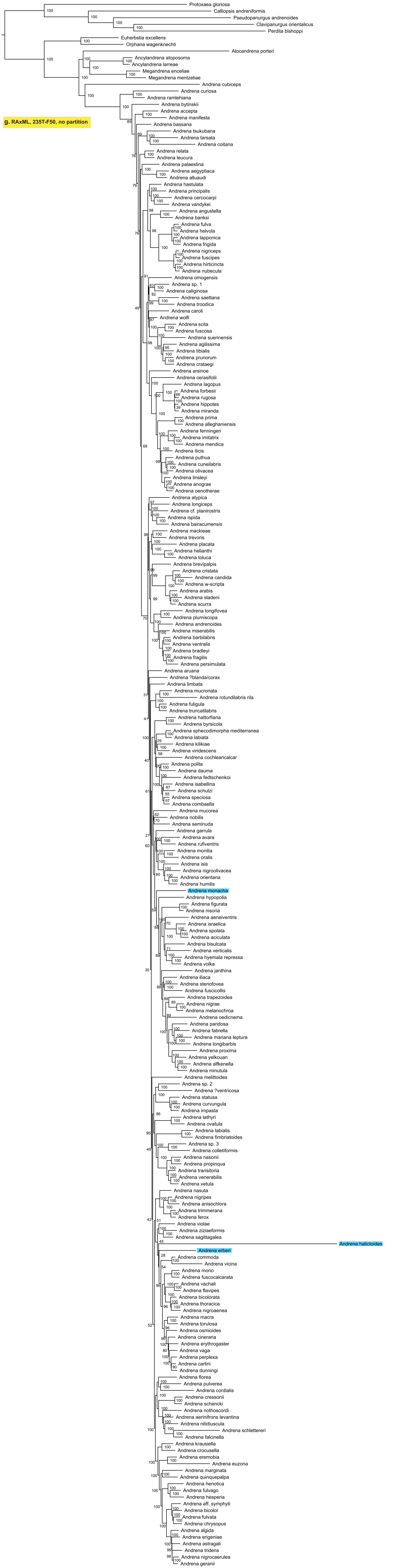

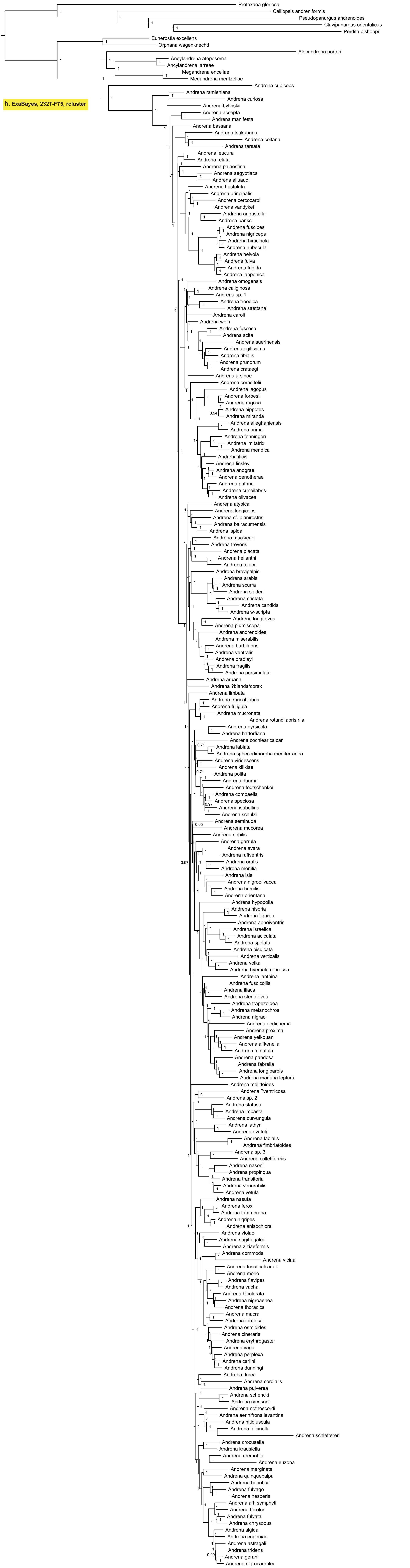

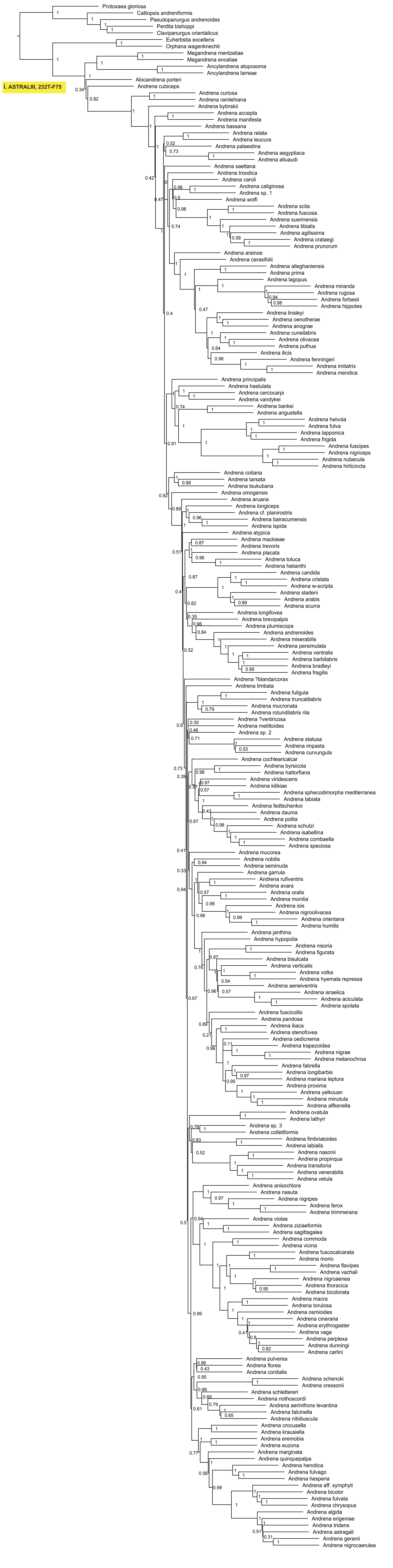

### Figure S2

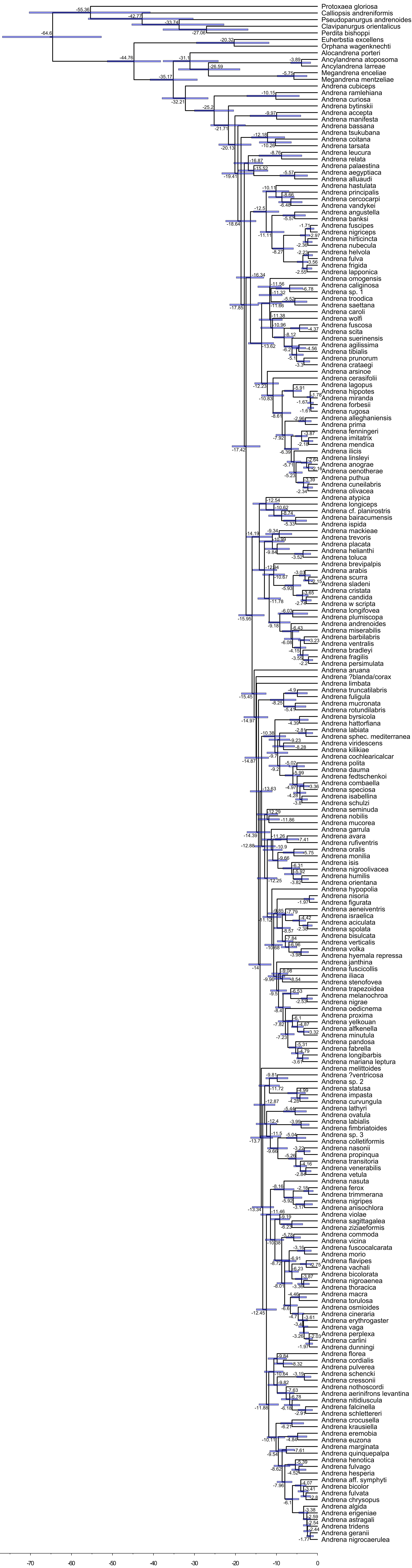

### Figure S3

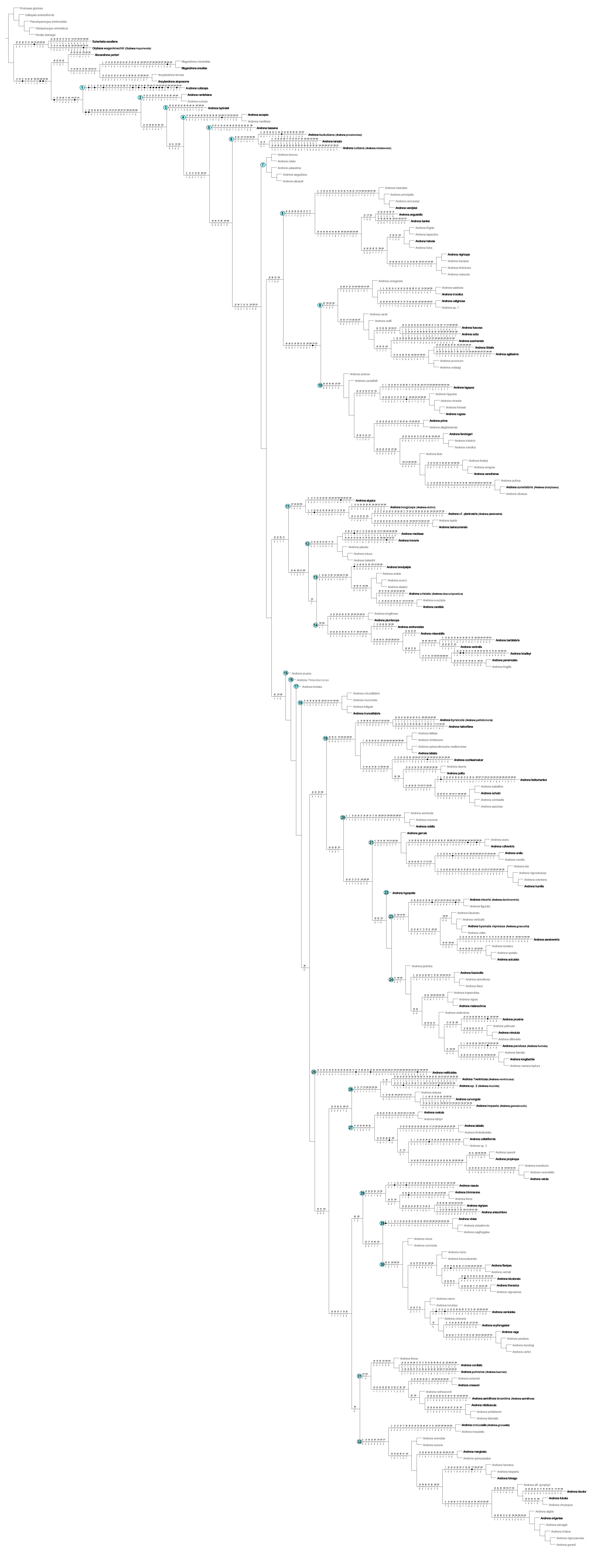

### Figure S4

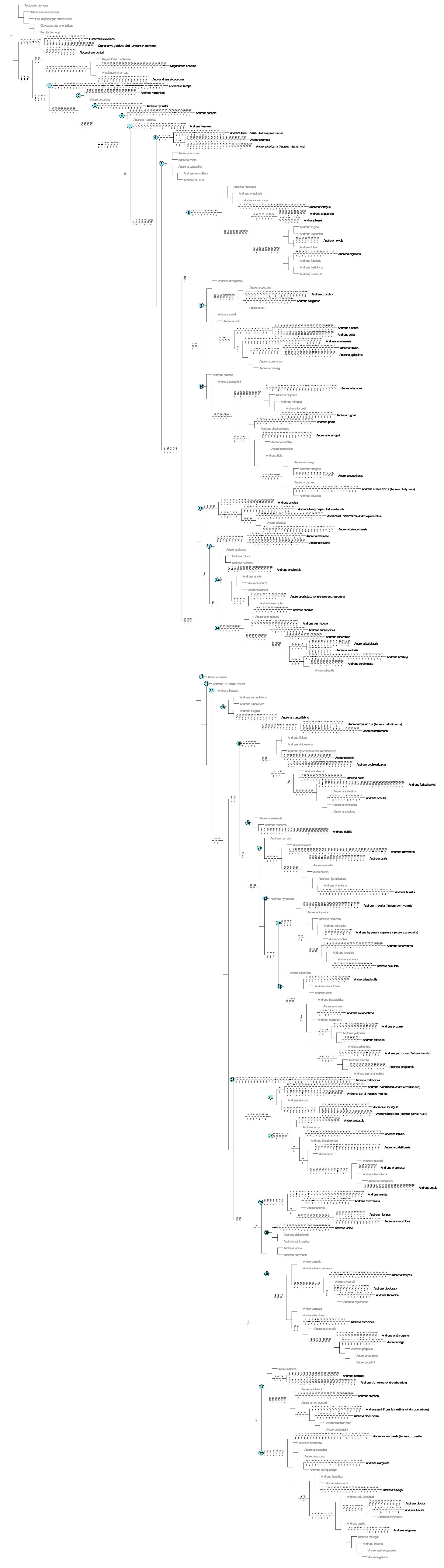
