## Supplementary material for "Molecular phylogeny and historical biogeography of andrenine bees (Hymenoptera: Andrenidae)": Table S3

**Table S3.** Ancestral ranges inferred by the Lagrange model for Andreninae nodes. Only nodes with alternative ranges within 2 log-likelihood units of the maximum for each node are shown. Ancestral ranges are presented in order of likelihood value (lnL). For each split, ranges on the left vs. right hand side correspond to upper vs. lower branches in Fig. 1. Clade terminology corresponds to Fig. 1.

| **node** | **split** | **lnL** | **rel. prob.** |
| --- | --- | --- | --- |
| Andreninae | [NT\|NA+NT] | -145.4 | 0.50 |
|  | [NT\|NA] | -145.9 | 0.29 |
|  | [NT\|NT] | -146.3 | 0.21 |
| Andrenini | [NA+NT\|NA] | -145.0 | 0.72 |
|  | [NA\|PA] | -146.7 | 0.14 |
| Andrena | [PA\|PA+NA] | -144.9 | 0.82 |
|  | [PA\|PA] | -146.7 | 0.14 |
| clades 2–32 | [PA\|PA+NA] | -144.9 | 0.85 |
|  | [PA\|PA] | -146.8 | 0.13 |
| clades 7–32 | [PA\|PA] | -144.8 | 0.87 |
|  | [PA\|PA+NA] | -146.8 | 0.13 |
| clades 8+9+10 | [PA\|PA] | -145.4 | 0.48 |
|  | [NA\|PA+NA] | -145.8 | 0.32 |
|  | [NA\|NA] | -147.0 | 0.10 |
| clade 8 | [NA\|PA+NA] | -145.3 | 0.52 |
|  | [NA\|NA] | -145.5 | 0.47 |
| Anchandrena+Archiadrena+Cnemidandrena+  Andrena (i) | [NA\|PA+NA] | -145.4 | 0.49 |
|  | [NA\|NA] | -145.5 | 0.46 |
| Cnemidandrena+Andrena (i) | [NA\|NA] | -145.9 | 0.30 |
|  | [PA+NA\|PA] | -146.0 | 0.27 |
|  | [PA\|PA+NA] | -146.7 | 0.13 |
|  | [NA\|PA+NA] | -146.7 | 0.13 |
|  | [PA+NA\|NA] | -146.9 | 0.12 |
|  | [PA\|PA] | -147.8 | 0.04 |
| Andrena (i) | [PA\|PA+NA] | -144.9 | 0.82 |
|  | [PA\|PA] | -146.4 | 0.18 |
| clades 9+10 | [PA\|PA] | -145.4 | 0.50 |
|  | [PA\|PA+NA] | -146.1 | 0.26 |
|  | [PA+NA\|PA] | -146.6 | 0.14 |
| clade 9 | [PA\|PA] | -145.2 | 0.62 |
|  | [PA+NA\|PA] | -145.7 | 0.38 |
| Habromelissa+Dactylandrena+Andrena sp. 1+ Troandrena | [PA\|PA] | -145.2 | 0.58 |
|  | [PA\|PA+NA] | -145.6 | 0.42 |
| Suandrena+Agandrena+Plastandrena (i)+ Plastandrena (ii) | [PA\|PA] | -145.0 | 0.73 |
|  | [PA\|PA+NA] | -146.0 | 0.27 |
| clade 10 | [PA\|PA] | -145.4 | 0.50 |
|  | [PA\|PA+NA] | -145.9 | 0.29 |
|  | [PA\|NA] | -146.2 | 0.22 |
| first internal node of clade 10 | [NA\|PA+NA] | -145.0 | 0.78 |
|  | [NA\|NA] | -146.2 | 0.21 |
| second internal node of clade 10 | [PA+NA\|NA] | -144.9 | 0.80 |
|  | [NA\|NA] | -146.4 | 0.19 |
| clades 11–32 | [PA\|PA] | -145.0 | 0.76 |
|  | [PA+NA\|PA] | -146.2 | 0.22 |
| Leucandrena (i)+Larandrena (ii)+Conandrena+ Gonandrena | [NA\|NA] | -145.1 | 0.69 |
|  | [PA+NA\|NA] | -146.0 | 0.27 |
| Simandrena (i)+Simandrena (ii)+Simandrena (iii)+ Ptilandrena (i) | [PA\|PA] | -145.0 | 0.71 |
|  | [PA+NA\|PA] | -146.0 | 0.28 |
| clades 28+29+30 | [PA\|PA] | -145.2 | 0.60 |
|  | [PA+NA\|NA] | -146.2 | 0.23 |
|  | [PA\|PA+NA] | -146.9 | 0.11 |
| clade 28 | [PA\|PA] | -145.2 | 0.59 |
|  | [PA\|PA+NA] | -145.6 | 0.41 |
| clades 29+30 | [NA\|PA+NA] | -145.0 | 0.74 |
|  | [NA\|NA] | -146.1 | 0.25 |
| clade 30 | [NA\|PA+NA] | -144.9 | 0.78 |
|  | [NA\|NA] | -146.4 | 0.18 |
| first internal split in clade 30 | [PA\|PA+NA] | -145.0 | 0.71 |
|  | [PA\|NA] | -146.1 | 0.25 |
| second internal split in clade 30 | [NA\|PA+NA] | -145.0 | 0.74 |
|  | [NA\|NA] | -146.1 | 0.25 |
| third internal split in clade 30 | [NA\|PA+NA] | -144.9 | 0.80 |
|  | [NA\|NA] | -146.4 | 0.19 |
| Poliandrena (v)+Cordandrena+Hesperandrena | [PA\|PA] | -144.8 | 0.87 |
|  | [PA\|PA+NA] | -146.8 | 0.13 |

rel. prob., relative probability; NT, Neotropic; NA, Nearctic; PA, Palaearctic
